# An unexpected contribution of lincRNA splicing to enhancer function

**DOI:** 10.1101/287706

**Authors:** Jennifer Y. Tan, Adriano Biasini, Robert S. Young, Ana C. Marques

## Abstract

Transcription is common at active mammalian enhancers sometimes giving rise to stable and unidirectionally transcribed enhancer-associated long intergenic noncoding RNAs (elincRNAs). ElincRNA expression is associated with changes in neighboring gene product abundance and local chromosomal topology, suggesting that transcription at these loci contributes to gene expression regulation in *cis*. Despite the lack of evidence supporting sequence-dependent functions for most elincRNAs, splicing of these transcripts is unexpectedly common. Whether elincRNA splicing is a mere consequence of their cognate enhancer activity or if it directly impacts enhancer-associated *cis*-regulation remains unanswered.

Here we show that elincRNAs are efficiently and rapidly spliced and that their processing rate is strongly associated with their cognate enhancer activity. This association is supported by: their enrichment in enhancer-specific chromatin signatures; elevated binding of co-transcriptional regulators, including CBP and p300; increased local intra-chromosomal DNA contacts; and strengthened *cis*-regulation on target gene expression. Using nucleotide polymorphisms at elincRNA splice sites, we found that elincRNA splicing enhances their transcription and directly impacts *cis*-regulatory function of their cognate enhancers. Importantly, up to 90% of human elincRNAs have nucleotide variants that are associated with both their splicing and the expression levels of their proximal genes.

Our results highlight an unexpected contribution of elincRNA splicing to enhancer function.

## INTRODUCTION

Enhancers are distal DNA elements that positively drive target gene expression (Banerji et al., 1981; Li et al., 2016; Moreau et al., 1981). These regulatory regions are DNase I hypersensitive, marked by histone 3 acetylation at lysine 27 (H3K27ac) and a high ratio of monomethylation versus trimethylation at histone 3 lysine 4 (H3K4me1 and H3K4me3, respectively). Together, these chromatin signatures are commonly used to annotate enhancers genome-wide (Hoffman et al., 2012). Most active enhancers are also transcribed (De Santa et al., 2010; Kim et al., 2010). Relative to non-transcribed enhancers, those that give rise to enhancer-associated transcripts are more strongly associated with enhancer-specific chromatin signatures (Wang et al., 2011) and display higher levels of reporter activity both *in vitro* (Wu et al., 2014; Young et al., 2017) and *in vivo* (Andersson et al., 2014), supporting the link between enhancer transcription and *cis*-regulatory function. While most enhancers bidirectionally transcribed short noncoding RNAs that are non-polyadenylated, unspliced and short-lived (eRNAs) (Kim et al., 2010), a subset of enhancers is transcribed only in one direction. In contrast to eRNAs, these enhancers produce polyadenylated noncoding transcripts that are relatively long, stable, and frequently spliced (Hon et al., 2017; Koch et al., 2011; Marques et al., 2013). We refer to these intergenic enhancer-associated transcripts as elincRNAs (Marques et al., 2013).

Enhancer transcription can increase local chromatin accessibility (Mousavi et al., 2013), modulate chromosomal interactions between cognate enhancer and target promoters (Lai et al., 2013) and regulate the load, pause and release of RNA Polymerase II (RNAPlI) (Maruyama et al., 2014; Schaukowitch et al., 2014), ultimately contributing to enhanced expression of neighboring protein-coding genes (Marques et al., 2013). Recently, we showed that elincRNAs preferentially locate at topologically associating domain boundaries (TADs) and their expression correlates with changes in local chromosomal architecture (Tan et al., 2017). While the association between elincRNA transcription and enhancer activity is relatively well established, whether the molecular mechanisms underlying their functions depend on their transcript sequences has not yet been equivocally demonstrated. Notably, consistent with the absence of nucleotide conservation at their exons (Marques et al., 2013), many elincRNA functions appear to rely on transcription alone (Alexanian et al., 2017; Hsieh et al., 2014; Lai et al., 2013; Li et al., 2013; Yoo et al., 2012).

A relatively large proportion of elincRNAs is not only stably transcribed but also undergoes splicing (Hon et al., 2017; Marques et al., 2013). Recently, the role of candidate elincRNA splicing in *cis*-gene expression regulation was demonstrated. For example, splicing of one lincRNA expressed in mouse embryonic stem cells (mESCs), Blustr, whose transcriptional start site initiates from an active enhancer (Mouse Encode Consortium et al., 2012), was shown to be sufficient to modulate the expression of its cognate protein-coding gene target in *cis* (Engreitz et al., 2016). Importantly, removing the splicing signal of another elincRNA, Haunt, by replacing its endogenous locus with its cDNA, could not rescue its *cis* regulatory function (Yin et al., 2015). These anecdotal evidence corroborating the contribution of splicing to elincRNA *cis*-regulatory functions, together with the lack of compelling evidence supporting a transcript-dependent role for most elincRNAs, raise important questions on the role, if any, of elincRNA splicing to enhancer function.

Interestingly, alongside its role in intron removal and appropriate exon assembly, splicing also contributes directly to other aspects of RNA metabolism, including transcription (Le Hir et al., 2003). For example, transcription of intronless transgenes in mice is at least 10 times less efficient than that of their intron-containing counterparts (Brinster et al., 1988). DNA elements embedded within introns have also been shown to contribute to transcriptional regulation (Sleckman et al., 1996) and components of the spliceosome can directly enhance RNAPII initiation (Kwek et al., 2002) and transcript elongation (Fong and Zhou, 2001).

To assess the contribution of elincRNA splicing to *cis* gene regulation, we investigated elincRNA splicing and its link to cognate enhancer function. Unexpectedly, we found that elincRNAs are as efficiently spliced as protein-coding genes and that their maturation associates with stronger enhancer activity. Finally, population analysis of human nucleotide variants that alter elincRNA splice sites and statistical genomics further revealed a direct role of splicing in the regulation of elincRNA transcription and cognate enhancer function.

## RESULTS

We considered all DNase I hypersensitive regions in mouse embryonic stem cell (mESC) overlapping transcribed intergenic mESC enhancers (Methods (Encode Project Consortium, 2012)). As expected, most enhancers are predominantly bidirectionally transcribed (Figure 1A) and producing eRNAs (Kim et al., 2010). Around 5% of transcribed enhancers are unidirectionally transcribed and give rise to enhancer-associated lincRNAs (elincRNAs, Table ST1). The transcription profile of elincRNAs (Figure 1B) resembles that of other mESC-expressed lincRNAs (oth-lincRNAs) (Figure 1C) and expressed protein-coding genes (Figure 1D).

**Figure 1.**
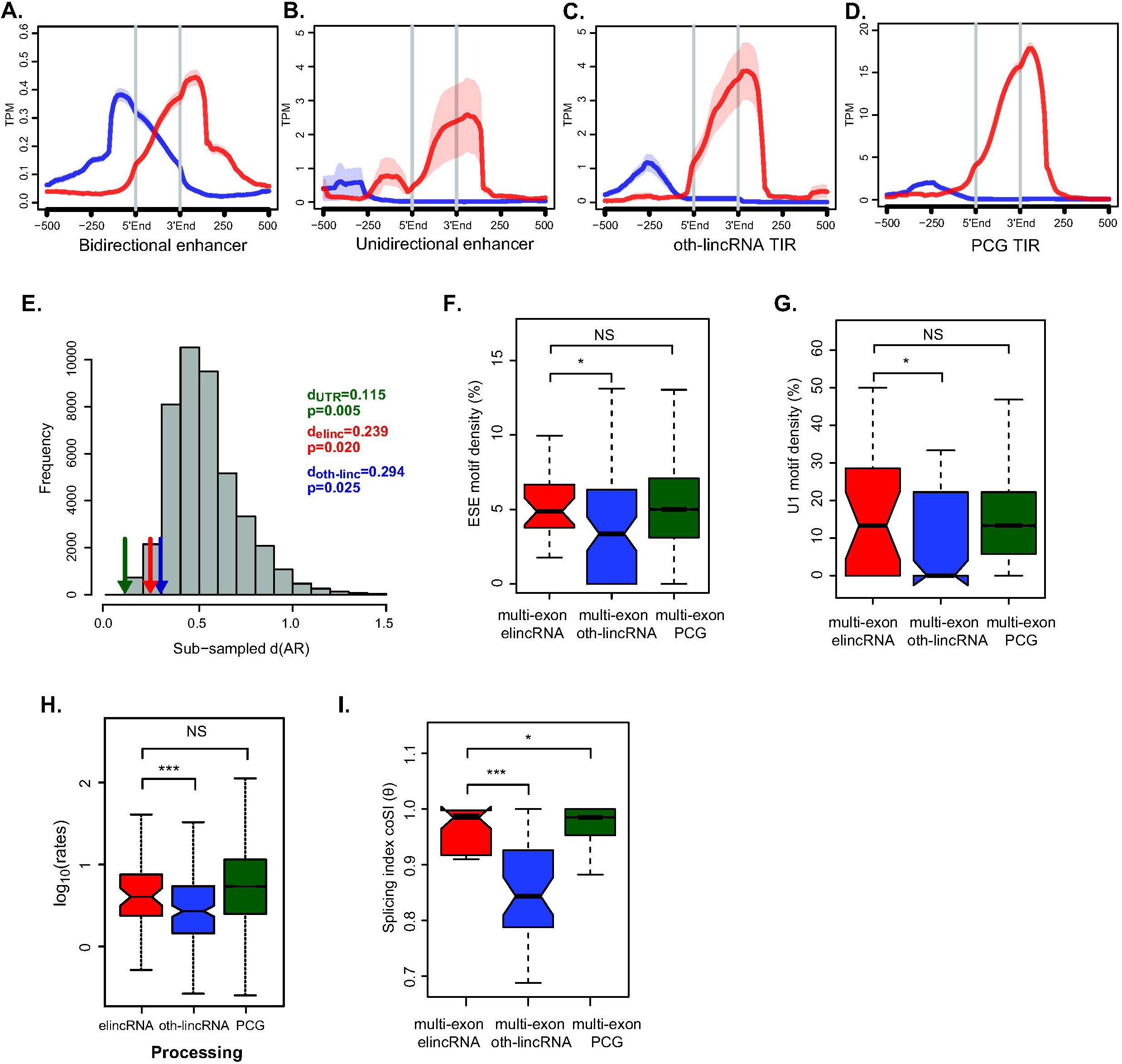
Rapid elincRNA splicing evolves under selection. Metagene plots of CAGE reads centered at transcription initiation regions (TIRs) of (A) eRNAs, (B) elincRNAs, (C) other mESC-expressed lincRNAs (oth-lincRNAs) and (D) protein-coding genes (PCGs). Sense (red) and antisense (blue) reads denote those that map to the same or opposite strand, respectively, as the direction of their cognate TIRs. (E) Pairwise substitution rate between mouse and human at splice sites of multi-exonic elincRNAs (red arrow), other expressed lincRNAs (blue arrow) and protein-coding gene UTRs (green arrow) relative to a background distribution built using 1,000 randomly subsampled sets of non-overlapping local ARs with matching GC-content and size. Distribution of the density of predicted (F) exonic splicing enhancers (ESEs) and (G) U1 spliceosome RNAs (snRNPs) within multi-exonic elincRNAs (red), other expressed lincRNAs (blue) and protein-coding genes (green). (H) Distribution of the average processing rates for elincRNAs (red), other expressed lincRNAs (blue) and protein-coding genes (green). (I) Distribution of the splicing index, coSI (θ) for multi-exonic elincRNAs (red), other expressed lincRNAs (blue) and protein-coding genes (green). Differences between groups were tested using a two-tailed Mann-Whitney *U* test. * *p* < 0.05; *** *p* < 0.001; NS *p* > 0.05.

### Rapid elincRNA splicing is associated with efficient transcription

Unlike eRNAs, elincRNAs are commonly spliced (44% are multi-exonic, median exon count=3) and their exons and introns display distinct GC contents, similar to protein-coding genes and other lincRNA loci (Figure S1A) (Haerty and Ponting, 2015; Schuler et al., 2014). Difference in GC content between intronic and exonic sequences is known to facilitate splice-site recognition and increase splicing efficiency (Amit et al., 2012). Supporting the biological relevance of elincRNA splicing, we found that selection has purged mutations at their splice sites (SS) (Figure 1E), and that their SS-flanking regions are enriched in splicing-associated elements, including exonic splicing enhancers (ESEs, Figure 1F) and U1 snRNPs (Figure 1G). Relative to other multi-exonic lincRNAs, elincRNAs splice sites also have a higher likelihood of being recognized by the splicing machinery (Figure S1B-C).

To assess whether the strong selective constraint at multi-exonic elincRNA splice sites and their enrichment in splicing motifs reflected efficient transcript splicing at these loci, we determined the transcriptome-wide rates of synthesis, splicing and degradation in mESCs. Towards this end, we performed 4-thiouridine (4sU) metabolic labelling of RNA followed by sequencing (Methods). Consistent with previous reports, while lincRNAs as a class were significantly less efficiently spliced than protein-coding genes (Mele et al., 2017; Mukherjee et al., 2017), we found that, relative to other expressed lincRNAs, elincRNA transcripts were 1.5-fold more rapidly processed (Figure 1H) and 14% higher proportion of their introns have undergone complete splicing (Figure 1I)(p<0.05, two-tailed Mann-Whitney U test, Figure 2A-B, Table ST2). Importantly, the splicing efficiency of elincRNAs is comparable to that of protein-coding genes (Figure 1H-I). No significant differences were found in the synthesis and degradation rates between elincRNAs and other expressed lincRNAs (p>0.16 two-tailed Mann-Whitney U test, Supplementary Figure S1D).

**Figure 2.**
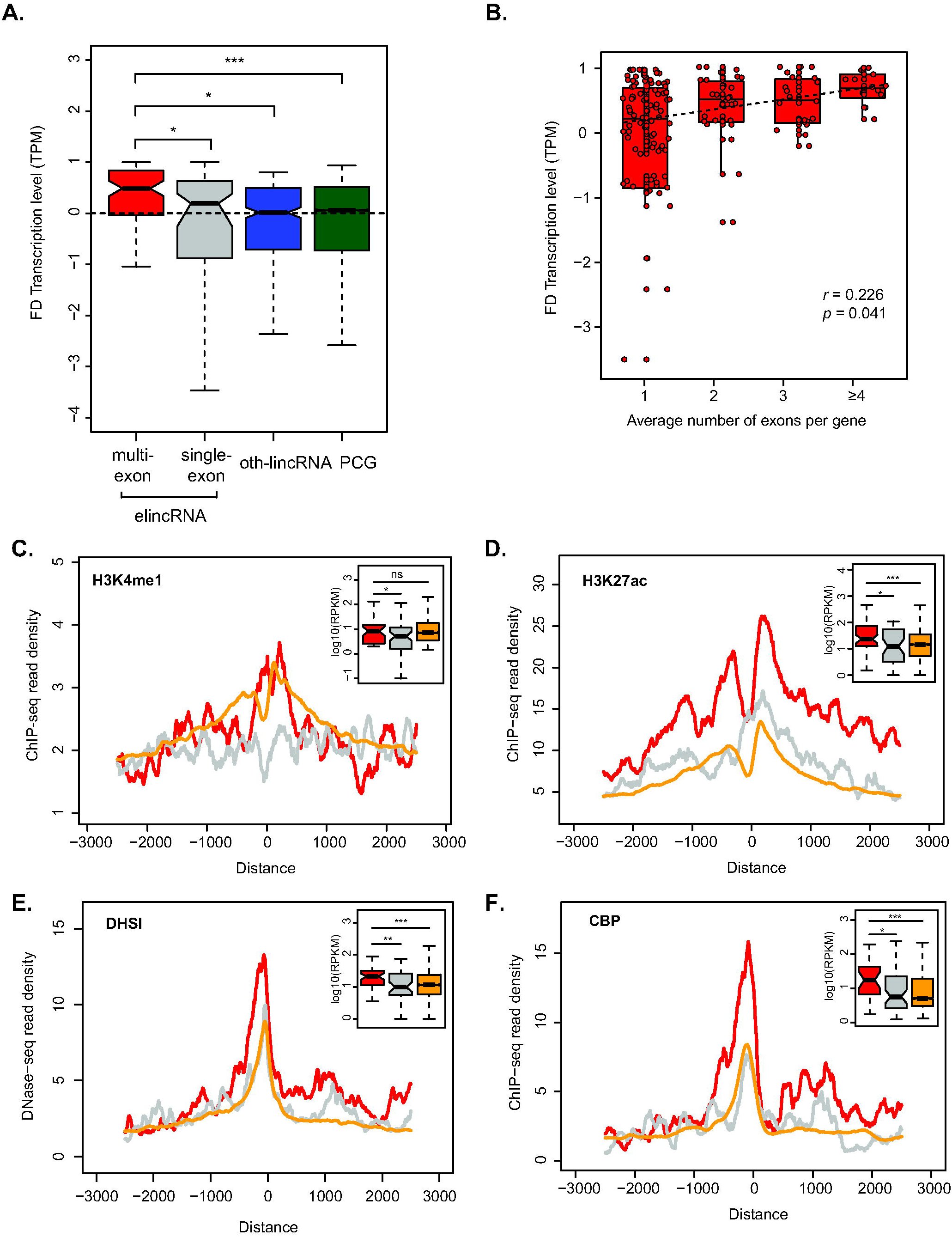
Multi-exonic elincRNAs are associated with higher enhancer activity. (A) Distribution of the fold difference (FD) in transcription (measured as CAGE TPM) of the most proximal gene to multi-exonic (red) and single-exonic (grey) elincRNAs, other mESC-expressed lincRNAs (oth-lincRNAs, blue) and protein-coding genes (PCGs, green) both expressed in a same stage of embryonic neurogenesis. Fold difference of neighboring genes is calculated between the two cellular stages across neuronal differentiation, where the expression level of their reference locus (elincRNA, oth-lincRNA, or PCG) is maximal and minimal. (B) Distribution of transcription FD for neighboring genes of elincRNAs with 1, 2, 3 or more than 4 exons (Spearman’s correlation). Metagene plots and distribution (figure insets) of (C) H3K4me1, (D) H3K27ac, (E) DNase I hypersensitive sites (DHSI) and (F) CBP ChIP-seq reads in mESCs at transcription initiation regions of multi-exonic (red) and single-exonic (grey) elincRNAs, and eRNAs (yellow). Differences between groups were tested using a two-tailed Mann-Whitney *U* test. * *p* < 0.05; *** *p* < 0.001; NS *p* > 0.05.

Surprisingly, we found multi-exonic elincRNA exons to have evolved neutrally (Figure S1E), suggesting the efficient splicing observed at these loci was not maintained through evolution to preserve the assembly of functionally-relevant sequence motifs within their primary transcript. Given the well-established coupling between splicing and transcription (Brinster et al., 1988; Le Hir et al., 2003), we questioned if splicing was instead associated with higher transcription of multi-exonic elincRNA loci. Consistent with this hypothesis, we found multi-exonic elincRNA transcripts were more rapidly synthesized compared to their single-exonic counterparts (Figure S1F). This higher transcriptional activity was supported by elevated levels of engaged RNA Polymerase II (RNAPII, Figure S1G) at their transcriptional initiation regions (TIRs) and lower RNAPII promoter-proximal stalling relative to other noncoding transcripts (Figure S1H, p<0.05, two-tailed Mann Whitney U test). Furthermore, relative to other non-spliced ncRNAs, multi-exonic elincRNA TIRs and gene bodies were enriched in phosphorylated Serine 5 (S5P) and Serine 2 (S2P) (Figure S1I-J), respectively, at RNAPII C-terminal domain, further supporting their high transcription initiation (Ho and Shuman, 1999), efficient transcription elongation and co-transcriptional splicing (Gu et al., 2013; Komarnitsky et al., 2000).

### Multi-exonic elincRNAs are associated with stronger enhancer activity

Next, we investigated whether the observed elincRNA splicing conservation and efficiency is linked to their cognate enhancer activity. During embryonic neurogenesis (Fraser et al., 2015), we found elincRNA transcription positively correlated with changes in neighboring protein-coding gene abundance (Figure S2A, Methods), similar to what was described previously (Marques et al., 2013). Strikingly, we found this association to be 2.5-fold stronger for multi-exonic elincRNAs than their single-exonic counterparts (p<0.05, two-tailed Mann-Whitney U test, Figure 2A). In contrast, no association was observed for other transcript classes, regardless of their splicing activity (Figure S2B). Expression changes in neighboring protein-coding gene abundance was correlated with the number of elincRNA exons (Figure 2B), suggesting that the amount of splicing occurring within multi-exonic elincRNAs may contribute to their *cis*-regulatory roles.

Consistent with their stronger association with neighboring protein-coding gene expression, chromatin signatures associated with high enhancer activity were found at enhancers that transcribe multi-exonic elincRNAs compared to those that give rise to either single-exonic elincRNAs or eRNAs. Specifically, multi-exonic elincRNA-producing enhancers were enriched for mono-methylation of Histone 3 Lysine 4 (H3K4me1, Figure 2C), acetylation of Histone 3 lysine 27 (H3K27ac, Figure 2D) and DNase I accessibility (DHSI, Figure 2E). Strikingly, using a hypothesis-free approach, we found that relative to their unspliced counterparts, TIRs of multi-exonic elincRNAs were significantly enriched (FDR<0.05) for transcription factor binding motifs required for the recruitment of the transcriptional co-activator CREB-binding protein (CBP) (Bedford et al., 2010), including Stat1, Egr1, Sp2, Smad3 and Klf5 (Table ST3). For a subset of the enriched CBP-recruiting transcription factors with available ChIP sequencing data in mESCs, and the CBP transcriptional co-activator, p300 (Merika et al., 1998), we found experimental support for their more frequent binding at multi-exonic elincRNAs’ TIRs (Figure 2F, S2C-E). Interestingly, direct binding of CBP to enhancer-associated RNAs was recently demonstrated to stimulate its histone acetylation activity and induce activation of target gene transcription (Bose et al., 2017). Our findings raise the possibility that spliced-elincRNAs are more likely to physically interact with CBP than are other enhancer-derived RNAs.

### Multi-exonic elincRNAs are specifically associated with changes in local chromosomal architecture

Since *cis*-regulatory interactions are known to be highly dependent on local chromosomal architecture, we examined whether the observed association between elincRNA splicing and enhanced neighboring gene expression was mediated through the modulation of their local chromosomal organization.

Analysis of their relative position within mESC topologically associating domains (TADs) revealed that strikingly, only multi-exonic elincRNA TIRs were significantly enriched at TAD boundaries and depleted at TAD centers (p<0.05, Figure 3A, Methods). This suggests that elincRNAs’ preferential location at TAD boundaries (Tan et al., 2017) is restricted to spliced elincRNAs. Chromosomal looping between enhancers and promoters occur frequently at TAD boundaries (Lupianez et al., 2015; Symmons et al., 2014). Importantly, we found multi-exonic elincRNA TIRs to be enriched at loop anchors relative to TIRs of all enhancer-derived transcripts (1.45-fold enrichment, FDR<0.05, permutation test, Methods), supporting their role in modulating enhancer-promoter interactions. This is further supported by the enriched binding of protein factors implicated in the establishment and modulation of chromosomal topology (Bonev and Cavalli, 2016) at multi-exonic elincRNA-producing enhancers, relative to their single exonic counterparts, including Ctcf (Figure S3A), subunits of the cohesin complex (Smc1 and Smc3), its cofactor Nipbl (Figure S3B-D), and the mediator complex (Med1 and Med3) (Figure S3E-F) in mESCs. Interestingly, enhancer-associated transcripts participates in enhancer-promoter looping by recruiting Cohesin or Mediator complexes to enhancer regions, which in turn stimulate cognate target gene transcription (Hsieh et al., 2014; Lai et al., 2013; Li et al., 2013).

**Figure 3.**
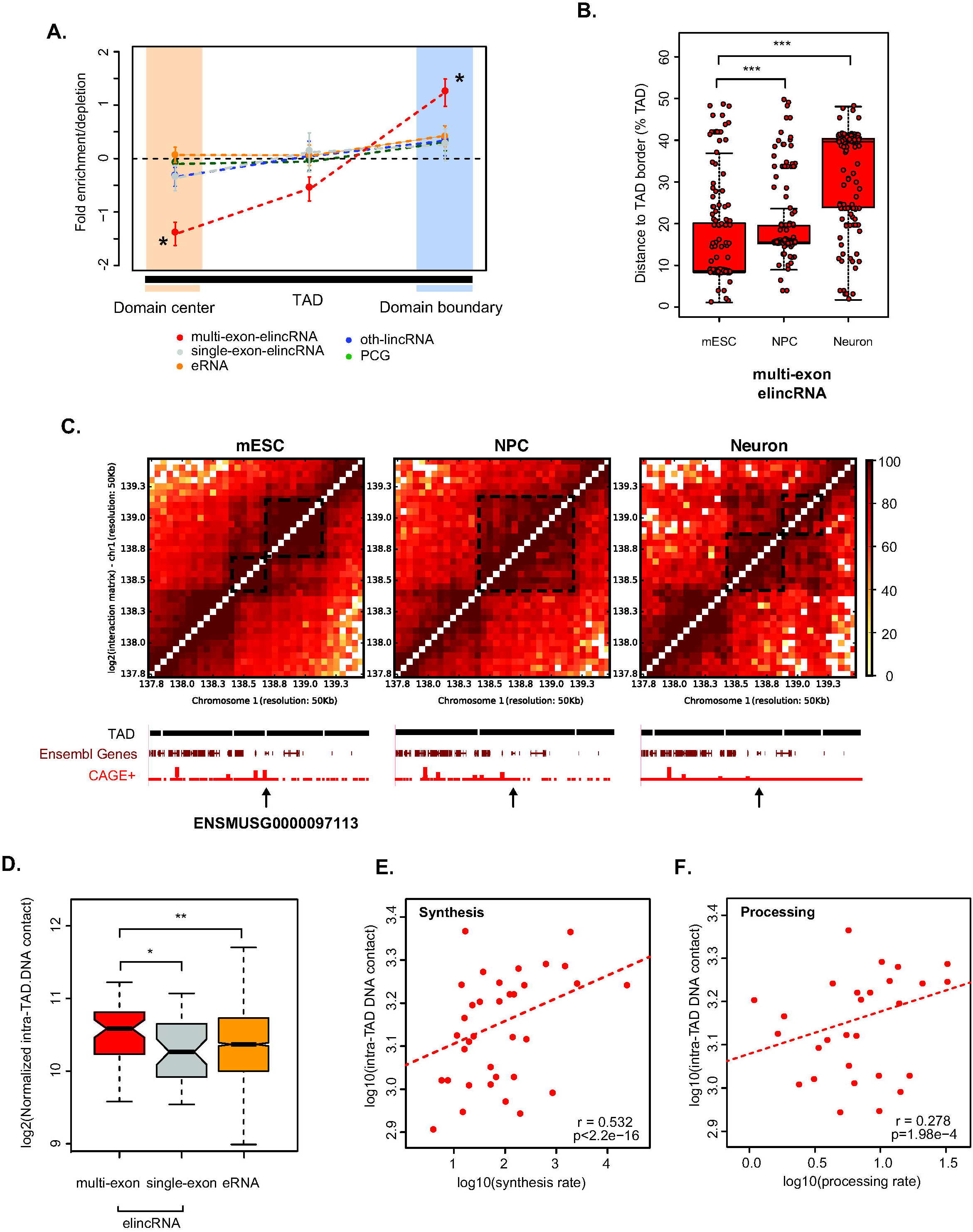
Multi-exonic elincRNAs are associated with modulation of local chromosomal architecture. (A) Fold enrichment or depletion of multi-exonic (red) and single-exonic (grey) elincRNAs, eRNAs (yellow), other expressed lincRNAs (blue) and protein-coding genes (green) at boundaries (light blue shaded area) and center (light yellow shaded areas) of TADs. Significant fold differences are denoted with *(*p*<0.05, permutation test) and standard deviation is shown with error bars. (B) Distribution of the distance between multi-exonic elincRNA transcription initiation site (red) to the nearest TAD border in mESCs, neuronal precursor cells (NPCs) and neurons. (C) Heatmap displaying the amount of chromosomal interactions, measured using HiC data, at regions surrounding one multi-exonic elincRNA (ENSMUSG0000097113) in mESC, NPC, and Neuron. Dotted black squares denote TAD, which is also represented by the black bars below the heatmap. Gene browser view of the corresponding region displaying Ensembl gene models (dark red lines) and CAGE read density (red lines) at each cell stage. (D) Distribution of the average amount of chromosomal contacts within mESC TADs that contain multi-exonic (red) and single-exonic (grey) elincRNAs and eRNAs (yellow). DNA-DNA contacts within multi-exonic elincRNA-containing mESC TADs (log10, Y axis) as a function of their respective (E) synthesis rate or (F) processing rate (log10, red points, Spearman’s correlation). Differences between groups were tested using a two-tailed Mann-Whitney *U* test. * *p* < 0.05; ** *p* < 0.01; *** *p* < 0.001; NS *p* > 0.05.

Consistent with the role of multi-exonic elincRNAs and their underlying enhancers in cell type-specific modulation of local chromosomal structure we found that whilst, on average, the location of single-exonic enhancer-derived lincRNAs and eRNAs remained relatively unchanged with respect to their nearest TAD border (Figure S4A), the distance between TAD borders and multi-exonic elincRNA TIRs increased upon cell differentiation (Figure 3B-C). Interestingly, multi-exonic elincRNA transcription is strongly correlated with the presence and maintenance of TAD boundaries across differentiation, supporting cell-type-specific functions of these enhancers (Figure S4B-C).

To assess the impact of multi-exonic elincRNA transcription on local chromosomal architecture, we next investigated the relationship between enhancer transcription and intra-TAD DNA contact density (Methods). We found that the frequency of DNA contacts within TADs that encompass multi-exonic elincRNA loci to be significantly higher than those containing other transcribed enhancers (p<0.05, two-tailed Mann-Whitney *U* test, Figure 3D, Methods). Furthermore, we found that the density of local chromosomal interactions correlated with the rate of transcription (Figure 3E) and processing (Figure 3F) of multi-exonic elincRNAs.

### Disruption of elincRNA splicing decreases target expression

The association between efficient elincRNA splicing and cognate enhancer activity can either reflect a direct role of noncoding RNA splicing in strengthening enhancer function or be a consequence of higher enhancer activity on transcriptional output. To distinguish between the two alternatives, we assessed the impact of single nucleotide polymorphisms (SNPs) (1000 Genomes Project Consortium, 2012) that disrupt elincRNA splice sites (Table ST4) on their putative target expression. We identified 38 variants disrupting splice donor acceptor sites within 37 elincRNAs. As expected, we found the percentage of full intron excision in individuals that carry variants that would disrupt splice junctions decreased by 6% relative to those carrying the reference canonical splice sites’ allele (GT-AG) (Figure 4A). Importantly and despite the relatively low fraction of affected splice junctions (average 13.5%), we found the relative abundance of elincRNAs was significantly decreased by 9.5% in individuals that carry variants that alter their splice donor acceptor sites (Figure 4B). Importantly, this natural mutational study revealed that decreases in elincRNA splicing was also associated with significant down-regulation of their putative protein-coding gene targets expression (p<8×10^-6^, two-tailed Mann-Whitney’s U test, Figure 4C) supporting a direct role of splicing in the modulation of enhancer function.

**Figure 4.**
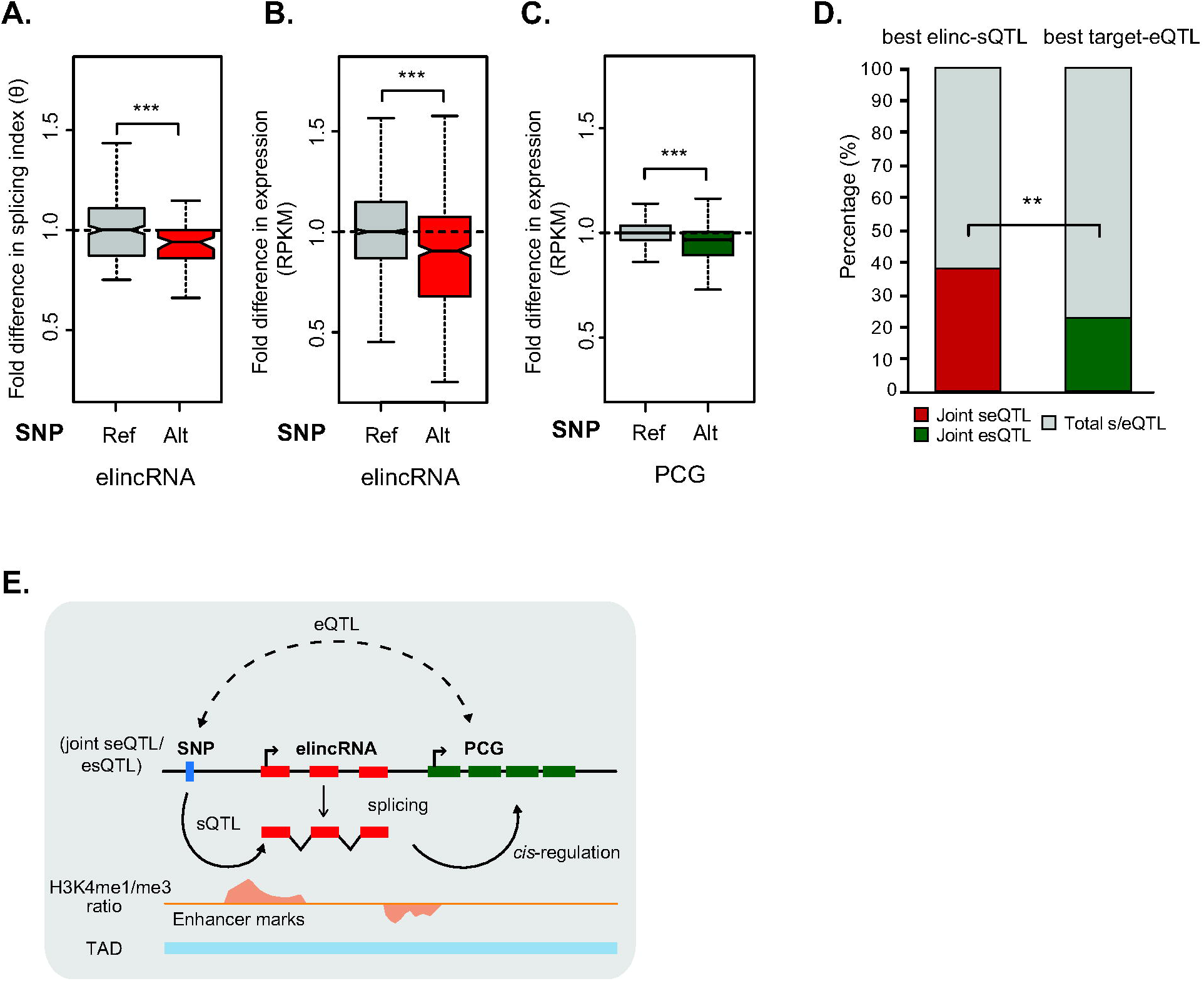
Impact of elincRNA splicing on *cis*-gene regulation in the human population. Distribution of the fold difference in the (A) splicing index and (B) expression levels (RPKM) of multi-exonic elincRNAs (red) and (C) expression levels (RPKM) of their putative target protein-coding genes (green) between individuals that carry the alternative allele (red, green) at elincRNA splice site SNPs relative to those that have the reference allele (grey). (D) The proportion of best elincRNA-sQTLs that are jointly associated with the expression levels (eQTLs) of their putative target protein-coding genes (joint seQTLs) (red) out of all elincRNA-sQTLs (grey) [Forward model] compared to the proportion of best target-eQTLs that are jointly associated with elincRNA-sQTLs (joint esQTLs, green) out of all target-eQTLs (grey) [Reverse model]. Differences between groups were tested using a two-tailed Mann-Whitney *U* or Fisher’s exact test. ** *p* < 0.01; *** *p* < 0.001. (E) Mediated through splicing of elincRNAs (red), genetic variants associated with elincRNA splicing (sQTLs) are likely to be indirectly associated with the expression level (eQTLs) of their putative *cis*-target genes (green). Spliced elincRNAs are preferentially located at topologically associating domain (TAD) boundary (blue bar), and transcription of these loci initiates from enhancer regions marked by high H3K4me1/me3 ratio (yellow), and likely strengthens enhancer activity to regulate neighboring target gene expression in *cis*.

To account for the relatively low number of variants overlapping splice sites and to further assess the impact of elincRNA splicing on *cis*-regulation, we identified SNPs associated with the amount of splicing at multi-exonic elincRNA loci (sQTLs, Methods) but not with their expression (eQTLs). When we estimated the proportion of elincRNA-sQTLs also associated with their putative target expression (joint seQTLs) (Figure S5A), we found that strikingly, nearly 90% (104/116) of multi-exonic elincRNAs with splicing-associated variants had at least one sQTLs that was also associated with their target expression (40% of all sQTLs, n=6197, Figure S5B).

Since sQTLs associated with the same locus are likely to be in high linkage disequilibrium, we obtained a conservative set of elincRNA splicing variants by considering for each elincRNA only the sQTL with the strongest association (best-sQTL). Of these, 43 (37%) were jointly associated with their target gene expression (Figure 4D). Remarkably, we found a significantly lower proportion of target protein-coding gene best-eQTLs (22.3%, 45/206, p<1X10^-2^, two-tailed Fisher’s exact test) to be also associated with elincRNA-sQTLs (joint esQTLs), signifying the impact of elincRNA splicing on nearby gene *cis*-regulation is significantly greater than what would be expected by chance given their local chromatin environment. Statistical mediation test that assesses whether the association between target gene expression and their eQTL variants was an indirect consequence of elincRNA splicing predicted that for 40% (17/43) of these elincRNA-seQTL-target triplets, elincRNA splicing was likely to be the mediating factor in target gene expression (FDR<0.05, Sobel’s test, Methods, Figure S5A, C, 4E). This is almost 20-fold higher than when the expression of target protein coding gene was predicted to mediate elincRNA splicing (1/43, Figure S5A, C).

## DISCUSSION

Most active enhancers are bidirectionally transcribed and produce short and unstable eRNAs (Andersson et al., 2014). A fraction of these transcribed enhancers produces unidirectionally transcribed transcripts (elincRNAs) that are sometimes spliced (Hon et al., 2017; Marques et al., 2013). Here we sought to understand if differences in the directionality and transcript structure of enhancer-associated transcription underlie differences in enhancer activity and function. We found that enhancers that produce elincRNAs, particularly those that undergo splicing, are more strongly associated with: typical epigenetic signatures of highly active enhancers; greater fold increase in putative *cis*-target expression; and the modulation of local chromosomal architecture. Given the paucity of evidence supporting a sequence-dependent mechanism for most elincRNAs and their poor exonic nucleotide conservation, unexpectedly, we found splicing of elincRNAs to be not only conserved during evolution but also highly efficient. Our population genomics analysis further supports a link between elincRNA splicing and local gene expression regulation.

It was recently shown that newly evolved transcriptional initiation sites are intrinsically bidirectional (Jin et al., 2017) and that the acquisition of splicing and polyadenylation signals can favor the preservation of the preferred transcription direction (Almada et al., 2013). Given the rapid turnover (Villar et al., 2015) and bidirectional transcription (Andersson et al., 2014) found at most mammalian enhancers, we questioned whether differences in enhancer transcription directionality and the splicing ability of their associated transcripts reflect differences in their evolutionary age. In mESCs, more than half (57%) of enhancers that produce elincRNAs have conserved chromatin signatures at their syntenic regions in human ESCs, a significantly higher proportion than those that produce eRNAs (23%, p<5×10^-13^, two-tailed Fisher’s exact test). Importantly, most of the conserved enhancers give rise to spliced elincRNAs (80%), consistent with their relative old evolutionary age.

We propose that enhancers are initially bidirectionally transcribed and over time, evolved features, including splicing, that strengthened their transcription. This proposal is consistent with the frequent birth of exons during mammalian evolution and evidence that novel exon-containing isoforms are more highly expressed (Merkin et al., 2015). Higher enhancer transcription may facilitate the binding of molecular factors, such as CBP, the Cohesin and Mediator complexes, at their cognate enhancers, which was recently shown to induce local chromosomal remodeling (Bose et al., 2017; Hsieh et al., 2014; Lai et al., 2013; Li et al., 2013), ultimately leading to stronger enhancer activity observed at these loci.

## METHODS

### Identification of enhancer-associated transcripts

We considered mESC ENCODE intergenic enhancers (Bogu et al., 2015) to be transcribed if they overlapped DNase I hypersensitive sites (Mouse Encode Consortium et al., 2012) and a CAGE cluster (Fraser et al., 2015) in the corresponding cell type (n=2217). We considered all mESC-expressed lincRNAs (Tan et al., 2015) and ENSEMBL annotated protein coding genes (version 70) with at least one CAGE read overlapping (by > 1 nucleotide) their first exon and a mESC CAGE cluster on the same strand. One hundred transcribed enhancers overlapped lincRNA CAGE clusters (Table ST1). The remaining CAGE clusters were transcription initiation regions (TIRs) associated with 13,143 protein-coding genes and 317 other mESC-expressed lincRNAs (oth-lincRNAs). Coordinates of ENCODE-predicted enhancer elements in human GM12878 lymphoblastoid cells (LCLs) (Encode Project Consortium, 2012) were obtained after excluding those found within the ENCODE Data Analysis Consortium Blacklisted Regions (Hoffman et al., 2013). LCL-expressed lincRNAs (as described in (Tan et al., 2017)), whose transcription initiation regions overlap intergenic LCL enhancers, as described previously, were considered as elincRNAs.

Metagene profiles of CAGE reads centered at mESC enhancers and gene TIRs were plotted using NGSplot (Shen et al., 2014). Sense and antisense reads denote those that map to the same or opposite strand, respectively, as the direction of their cognate CAGE clusters. For eRNAs, direction is defined as the direction with the highest number of CAGE clusters. In cases of equal CAGE clusters on either direction, enhancer direction is randomly assigned.

### GC composition

Only mESC genes with multi-exonic transcripts (2 or more exons) were considered for this analysis. We computed GC content separately for the first and all remaining exons, as well as the introns, for each gene and their flanking intergenic sequences of the same length, after excluding the 500 nucleotides immediately adjacent to annotations, as previously described (Haerty and Ponting, 2015).

### Identification of splicing-associated motifs

We predicted the density of mouse exonic splicing enhancer (ESE) motifs (identified in (Fairbrother et al., 2002)) within mESC transcripts, as described previously (Haerty and Ponting, 2015). Exonic nucleotides (50 nt) flanking splice sites (SS) of internal transcript exons (> 100 nt) were considered in the analysis, after masking the 5 nt immediately adjacent to SS to avoid splice site-associated nucleotide composition bias (Fairbrother et al., 2002; Yeo and Burge, 2004). Canonical U1 sites (GGUAAG, GGUGAG, GUGAGU) adjacent to 5’ splice sites (3 exonic nt and 6 intronic nt flanking the 5’ SS) were predicted as previously described (Almada et al., 2013). FIMO (Grant et al., 2011) was used to search for perfect hexamer matches within these sequences. For each exon, we estimated the splice site strength using MaxENT (Yeo and Burge, 2004). SS scores were calculated using the −3 exonic nt to +6 intronic nt and −20 intronic nt to +3 exonic nt flanking the 5’ SS and 3’ SS, respectively.

### Evolutionary constraint analysis

Syntenic regions of mESC (mm9) genetic elements in human (hg19) were determined using LiftOver with parameters: -minMatch=0.2, -minBlocks=0.01 (Meyer et al., 2013). Regions within the ENCODE Data Analysis Consortium Blacklisted Regions (Hoffman et al., 2013) were excluded from this analysis.

To assess selective constraints, first, pairwise alignment of exons that belong to the same gene were concatenated and all splice site dinucleotides of the same gene biotype were concatenated. Next, their pairwise nucleotide substitution rates were estimated between mouse and human using BASEML from the PAML package [REV substitution model (Yang, 1997)]. Only sequences longer than 100 nt were considered in the analyses. Significance of nucleotide constraint was estimated by comparing the substitution rate of the region of interest to that observed for 1000 randomly simulated sets of non-overlapping adjacent (within 1 Mb) ancestral repeats (ARs) with matching GC-content and size between mouse and human.

### 4sU metabolic labelling of mESCs and RNA extraction

Mouse DTCM23/49 XY embryonic stem cell lines (mESCs) were cultured at 37°C with 5% CO_2_ in Knockout Dulbecco’s Modified Eagle Medium (D-MEM, Thermo Fisher, #10829-018) supplemented with 15% fetal bovine serum (FBS, Thermo Fisher, #16000-044), 1% antibiotic penicillin/streptomycin (Thermo Fisher, 15070063), 0.01% recombinant mouse LIF protein (Merck, #ESG1107) and 0.06 mM 2-Mercaptoethanol (Thermo Fisher, #31350-010), on 0.1% gelatin-coated cell culture dishes. When confluent, culture was divided in two and passaged 8 times. Five million mESCs of two biological replicates were seeded and allowed to grow to 70-80% confluency (approximately 1 day). RNA was labeled with 4sU (Sigma, T4509) and nascent RNA was isolated following the general procedure as previously described (Dolken et al., 2008). Specifically, 4sU was added to the growth medium (final concentration of 200 µM) and the cells were incubated at 37 °C for 15, 30 or 60 minutes. Plates were washed once with 1X PBS and RNA was extracted using Trizol (Thermo Fisher, #15596-026). 100 µg of extracted RNA was incubated for 2 h at room temperature with rotation in 1/10 volume of 10X biotinylation buffer (Tris-HCl pH 7.4, 10 mM EDTA) and 2/10 volume of biotin-HPDP (1mg/ml in Dimethylformamide (Thermo Fisher, #21341)). RNA was extracted using phenol:chloroform:isoamyl alcohol (Sigma, P3803-400ML). Equal volume of biotinylated RNA and pre-washed Dynabeads^TM^ MyOne^TM^ Streptavidin T1 beads (Thermo Fischer, #65601) was added to 2X B&W Buffer (10 mM Tris-HCl pH7.5, 1mM EDTA, 2M NaCl (Thermo Fisher, #65601)) and incubated at room temperature for 15 minutes under rotation. The beads were then separated from the mixture using DynaMag^TM^-2 Magnet (Thermo Fisher, #12321D). After removing the supernatant, beads were washed with 1X B&W three times. Biotinylated RNA was recovered from the supernatant after 1 minute of incubation with RTL buffer (RNeasy kit, Qiagen, #74104) and purified using the RNeasy kit according to manufacturer’s instructions.

### RNA sequencing, mapping, and quantification of metabolic rates

Total RNA libraries were prepared from 10 ng of DNase-treated total and newly transcribed RNA using Ovation^®^ RNA-Seq and sequenced on Illumina HiSeq 2500 (average of fifty million reads per library).

Hundred nucleotides long single-end stranded reads were first mapped to mouse ribosomal RNA sequences with STAR v2.5.0 (Dobin et al., 2013). Reads that do not map to ribosomal RNA were then aligned to intronic and exonic sequences using STAR and quantified using RSEM (Li and Dewey, 2011). Rates of synthesis, processing and degradation were independently inferred using biological duplicates at each labeling points using the INSPEcT Bioconductor package v1.8.0 (de Pretis et al., 2015). Biotype differences in the average rate across the 3 labeling times were used in the analyses (Table ST2). The raw sequencing data is available on the NCBI Gene Expression Omnibus (GEO) under accession number GSE111951.

### Splicing efficiency

The efficiency of splicing was assessed by estimating the fraction of transcripts for each gene where its introns were fully excised using bam2ssj (Pervouchine et al., 2013). The splicing index, coSI (θ), represents the ratio of total RNA-seq reads spanning exon-exon splice junctions (excised intron) over those that overlap exon-intron junctions (incomplete excision) (Tilgner et al., 2012).

### Metagene analysis of binding enrichment at elincRNAs

Enrichment of histone modifications, transcription factor binding, and gene expression levels were assessed using publically available mESC ChIP-seq and RNA-seq data sets. Downloaded data sets are listed in Table ST5.

For all downloaded data sets, adaptor sequences were first removed from sequencing reads with trimmomatic (version 0.33) (Bolger et al., 2014) and then aligned to the mouse reference genome (mm9) using HISAT2 (version 2.0.2) (Kim et al., 2015).

Metagene profiles of sequencing reads centered at gene TIRs were visualized using HOMER v4.7 (Heinz et al., 2010).

### RNAPII stalling

Distribution of RNAPII across the gene TIR and body, commonly used as an indicator of promoter-proximal RNAPII stalling and efficient transcription elongation, was estimated by calculating the travelling ratio. Using mESC RNAPII ChIP-seq data (Brookes et al., 2012). The travelling ratio represents relative read density at gene TIRs divided by that across the gene body (Reppas et al., 2006).

### Enhancer activity across embryonic neurogenesis

Level of gene transcription initiation (CAGE-based TPM at TIRs) at each of the three stages of neuronal differentiation (mESC to NPC to neuron) were downloaded from (Fraser et al., 2015)). Each locus was paired with its genomically-closest protein-coding gene, considered here as its putative *cis*-target. Only pairs where both loci were expressed in at least one embryonic neurogenesis stage were considered. For each gene, the two stages where the locus of interest was most highly or lowly expressed were determined and used to calculate the fold difference between the expression difference of its putative *cis*-target, as described previously (Marques et al., 2013).

### Prediction of enriched transcription factor motifs at mESC enhancers

We predicted DNA motifs for transcription factors enriched at multi-exonic elincRNA TIRs (+/-500 bp from the center of TIRs) relative to those that transcribe single-exonic elincRNAs and eRNAs. Enrichment of motifs of at least 8mer were predicted using FIMO (Grant et al., 2011). Enriched motifs matching to known transcription factor binding sites (JASPAR 2016 (Mathelier et al., 2016)) were predicted using TOMTOM (Gupta et al., 2007) with default parameters.

### Analysis of preferential location and chromosomal contact within topologically associating domains

mESC topologically associating domains (TADs) (Fraser et al., 2015) were divided into 3 equal size segments. Enrichment or depletion of enhancer-associated transcripts was estimated for each TAD region, relative to the expectation, using the Genome Association Tester (GAT) (Heger et al., 2013). Specifically, TAD positional enrichment was compared to a null distribution obtained by randomly sampling 10,000 times (with replacement) segments of the same length and matching GC content as the tested loci within mappable intergenic regions of TADs (as predicted by ENCODE (Hoffman et al., 2013)). To control for potential confounding variables that correlate with GC content, such as gene density, the genome was divided into segments of 10 Kb and assigned to eight isochore bins in the enrichment analysis. The frequency of chromosomal interactions within TADs was calculated using mESCs Hi-C contact matrices (Fraser et al., 2015), as previously described (Tan et al., 2017).

### Mapping of molecular quantitative trait loci (QTLs)

Expression values (RPKM) of multi-exonic elincRNAs and protein-coding genes in EBV-transformed LCLs derived from 373 individuals of European descent (CEU, GBR, FIN and TSI) were quantified (as described in (Tan et al., 2017)). The corresponding processed genotypes were downloaded from EBI ArrayExpress (accession E-GEUV-1) (Lappalainen et al., 2013). Quantification of splicing events was estimated using LeafCutter (Li et al., 2018). Single nucleotide polymorphisms (SNPs) located within the same TAD as the genes of interest were tested for association with elincRNA splicing (sQTLs) and with expression levels (eQTLs) of elincRNAs and protein-coding genes. Only SNPs with minor allele frequency (MAF) greater than 5% were considered in the QTL analyses. sQTLs and eQTLs were estimated using FastQTL v2.184 (Ongen et al., 2016). To assess the significance of the correlation globally, we permuted the splicing or expression levels of each gene 1000 times and considered only sQTLs or eQTLs with an absolute regression coefficient greater than 95% of all permuted values to be significant. We further performed Benjamini-Hochberg multiple testing correction to estimate FDR (<5%) for all SNPs within the same TAD. Putative protein-coding gene targets of multi-exonic elincRNAs were predicted as those that reside within the same TADs and whose expression levels were associated to the same SNP variant as the expression of the elincRNAs.

### Impact of genetic variation at elincRNA splice sites on cis-gene expression

We considered all SNPs located at elincRNA splice sites and estimated the fold difference in elincRNA splicing and steady state abundance as well as in their putative target expression between individuals that carry the reference or alternative alleles of these variants (Table ST4).

### Causality inference between elincRNA splicing and nearby protein-coding gene expression

To infer the causal relationship between elincRNA splicing and nearby gene expression, we focused on QTLs that are associated with both splicing of elincRNAs and their putative target gene abundance. First, we estimated the proportion of elincRNA sQTLs that are jointly associated with expression of their nearby protein-coding genes (seQTLs) and compared this to the proportion of protein-coding gene eQTLs also associated with their proximal elincRNA splicing (esQTLs) using a two-tailed Fisher’s exact test. elincRNA sQTL variants that were also associated with elincRNA expression level or splicing of their putative *cis*-target genes were excluded from the analysis (n=14,575 out of 30,183).

We defined the best-sQTL for each elincRNA as the variant with the highest absolute regression slope value. For all best-elincRNA-sQTLs that were jointly associated with nearby gene expression (seQTLs), we performed a Sobel’s test of mediation (Sobel, 1982) on all triplets of seQTL – elincRNA splicing – target gene expression by independently testing two models: (1) the causal model with elincRNA splicing as the molecular mediator of gene expression; and (2) the reactive model where gene expression mediates elincRNA splicing. Sobel’s test was implemented using the powerMediation R package (Weiliang, 2018).

### Statistical tests

All statistical analyses were performed using the R software environment for statistical computing and graphics (R Development Core Team, 2008).

## Acknowledgements

We thank Chris P. Ponting and members of the Marques group for valuable comments and discussion. We thank Zoltán Kutalik and Olivier Delaneau for discussion on population genomics analysis; Francesco Nicassio and Matteo Marzi for help establishing 4sU labeling; and Mattia Pelizzola and Stefano de Pretis for advice on RNA metabolic rate inference. This work is funded by the Swiss National Science Foundation grant (PP00P3_150667 to ACM) and the NCCR RNA & disease. R.S.Y. acknowledges the support of the UK Medical Research Council (U127597124) and the Medical Research Foundation.

## Authors’ contributions

JYT and ACM designed the study. JYT, AB and RSY performed analyses. JYT and ACM conceived methods and discussed the results. ACM supervised the analysis. JYT and ACM wrote the manuscript. All authors approved the manuscript.

