## Supplementary Materials for "An unexpected contribution of lincRNA splicing to enhancer function"

| elincRNA_id | elincRNA_coordinates (mm9) | Avg_number_exons |
|---|---|---|
| ENSMUSG00000097188 | chr19:8797986-8800932 (+) | 4 |
| XLOC_006239 | chr11:117837516-117842138 (+) | 2 |
| ENSMUSG00000085412 | chr6:52052933-52069727 (+) | 14 |
| XLOC_025721 | chr3:121072505-121091611 (+) | 3 |
| ENSMUSG00000086617 | chr11:101471928-101477270 (+) | 3 |
| ENSMUSG00000075416 | chr2:30568667-30575577 (-) | 5 |
| XLOC_008541 | chr12:56608653-56659015 (+) | 3 |
| XLOC_011005 | chr13:114525859-114530002 (+) | 2 |
| ENSMUSG00000087523 | chr11:70468243-70469995 (-) | 2 |
| XLOC_012717 | chr14:65791169-65804492 (+) | 3 |
| ENSMUSG00000092274 | chr19:5824708-5845817 (-) | 4 |
| XLOC_000149 | chr1:30931864-30943540 (+) | 2 |
| XLOC_010895 | chr13:98062138-98066681 (+) | 2 |
| ENSMUSG00000085992 | chr11:95084604-95086946 (-) | 2 |
| XLOC_033652 | chr6:125382586-125410109 (+) | 6 |
| ENSMUSG00000097052 | chr2:26492697-26495764 (-) | 4 |
| ENSMUSG00000092341 | chr19:5795690-5802672 (-) | 2 |
| ENSMUSG00000097107 | chr10:21550305-21564657 (-) | 5 |
| XLOC_000719 | chr1:121202191-121217220 (+) | 4 |
| linc1260 | chr11:33434734-33444682 (-) | 3 |
| ENSMUSG00000056738 | chr2:129044705-129061340 (+) | 5 |
| ENSMUSG00000086290 | chr4:131864593-131866939 (+) | 5 |
| ENSMUSG00000097173 | chr11:22500338-22510882 (-) | 2 |
| ENSMUSG00000097050 | chr15:102100994-102104552 (+) | 2 |
| XLOC_035789 | chr7:80997054-81045284 (+) | 2 |
| ENSMUSG00000086152 | chr2:162948342-162969695 (+) | 3 |
| ENSMUSG00000086003 | chr11:93994776-94017090 (+) | 6 |
| XLOC_027549 | chr4:106532199-106534136 (+) | 2 |
| XLOC_003079 | chr10:72097074-72104866 (+) | 3 |
| ENSMUSG00000085156 | chr11:6425594-6428763 (-) | 4 |
| linc1557 | chr6:125382625-125408897 (+) | 4 |
| ENSMUSG00000097848 | chr13:99870657-99877535 (+) | 2 |
| XLOC_003300 | chr10:82806952-82814855 (+) | 2 |
| ENSMUSG00000074813 | chr2:128021729-128255085 (-) | 4 |
| ENSMUSG00000084829 | chr2:171018769-171096850 (+) | 5 |
| XLOC_012856 | chr14:99738198-99750535 (+) | 2 |
| XLOC_033363 | chr6:88102389-88104966 (+) | 2 |
| XLOC_027498 | chr4:99316961-99318579 (+) | 2 |
| ENSMUSG00000097113 | chr1:138567315-138587381 (+) | 5 |

| | | |
|---|---|---|
| ENSMUSG00000097697 | chr7:74929611-74948382 (-) | 3 |
| ENSMUSG00000097514 | chr2:170017949-170019553 (-) | 2 |
| XLOC_028145 | chr4:141127806-141131481 (+) | 2 |
| ENSMUSG00000086841 | chr11:62416379-62418309 (+) | 5 |
| XLOC_030498 | chr5:74337633-74340530 (+) | 2 |
| XLOC_016133 | chr16:84772549-84774033 (+) | 1 |
| XLOC_021148 | chr19:7335517-7336313 (-) | 1 |
| linc1418 | chr1:121202185-121202284 (+) | 1 |
| XLOC_000833 | chr1:135564142-135566485 (+) | 1 |
| XLOC_033583 | chr6:122315739-122316241 (+) | 1 |
| XLOC_000564 | chr1:88205133-88206690 (+) | 1 |
| XLOC_015105 | chr15:76888265-76889365 (-) | 1 |
| XLOC_015912 | chr16:32303909-32305082 (+) | 1 |
| XLOC_026247 | chr3:75457250-75459952 (-) | 1 |
| XLOC_025338 | chr3:88377003-88377402 (+) | 1 |
| XLOC_019554 | chr18:56837020-56837575 (+) | 1 |
| XLOC_037354 | chr7:80761834-80762820 (-) | 1 |
| XLOC_006215 | chr11:116953308-116954265 (+) | 1 |
| XLOC_025057 | chr3:35619769-35621125 (+) | 1 |
| XLOC_009927 | chr12:111187031-111188740 (-) | 1 |
| XLOC_000820 | chr1:135026005-135026936 (+) | 1 |
| XLOC_000291 | chr1:51803846-51804451 (+) | 1 |
| XLOC_018309 | chr17:24168770-24169322 (-) | 1 |
| XLOC_011274 | chr13:37531369-37531964 (-) | 1 |
| XLOC_004022 | chr10:69829774-69832071 (-) | 1 |
| XLOC_025507 | chr3:95455389-95456295 (+) | 1 |
| XLOC_022987 | chr2:165793743-165795294 (+) | 1 |
| XLOC_002128 | chr1:135566621-135568318 (-) | 1 |
| XLOC_014247 | chr15:77736036-77737127 (+) | 1 |
| XLOC_015658 | chr16:13749111-13750006 (+) | 1 |
| XLOC_010097 | chr13:17791134-17794983 (+) | 1 |
| XLOC_027921 | chr4:130174495-130175460 (+) | 1 |
| XLOC_040397 | chr9:44227398-44227945 (+) | 1 |
| XLOC_021469 | chr19:44354830-44355301 (-) | 1 |
| XLOC_038724 | chr8:86713767-86715539 (+) | 1 |
| XLOC_022429 | chr2:116798439-116799348 (+) | 1 |
| XLOC_020126 | chr18:56837013-56837936 (-) | 1 |
| XLOC_032353 | chr5:117852504-117855192 (-) | 1 |
| XLOC_033043 | chr6:39387892-39388140 (+) | 1 |
| XLOC_004758 | chr11:20037249-20039521 (+) | 1 |
| XLOC_039274 | chr8:32178049-32178300 (-) | 1 |
| XLOC_022829 | chr2:154252339-154254200 (+) | 1 |

| | | |
|---|---|---|
| XLOC_007295 | chr11:79787120-79789316 (-) | 1 |
| XLOC_017563 | chr17:37129288-37131980 (+) | 1 |
| XLOC_016817 | chr16:84772678-84774019 (-) | 1 |
| XLOC_036482 | chr7:3201216-3202855 (-) | 1 |
| XLOC_018391 | chr17:27176142-27177014 (-) | 1 |
| XLOC_029617 | chr4:140838533-140840319 (-) | 1 |
| XLOC_035137 | chr7:19510761-19511157 (+) | 1 |
| XLOC_007945 | chr11:116294651-116295113 (-) | 1 |
| XLOC_021281 | chr19:23161475-23162965 (-) | 1 |
| XLOC_025506 | chr3:95454805-95455230 (+) | 1 |
| XLOC_002753 | chr10:21546359-21547385 (+) | 1 |
| XLOC_015918 | chr16:32324193-32324738 (+) | 1 |
| XLOC_027890 | chr4:129059633-129060402 (+) | 1 |
| XLOC_036484 | chr7:3209749-3212199 (-) | 1 |
| XLOC_016350 | chr16:13749559-13750120 (-) | 1 |
| XLOC_017995 | chr17:88225276-88229061 (+) | 1 |
| XLOC_040760 | chr9:78351270-78351683 (+) | 1 |
| XLOC_023798 | chr2:71704980-71705877 (-) | 1 |
| XLOC_032351 | chr5:117849082-117849389 (-) | 1 |
