## Supplementary Materials for "An unexpected contribution of lincRNA splicing to enhancer function"

### Supplementary Table ST3. Enrichment of transcription factor binding motifs at elincRNA initiation sites

| | TF name | p-value | E-value | q-value | Motif |
|---|---|---|---|---|---|
| | Esr2 | 2.17E-05 | 3.11E-02 | 6.23E-02 | 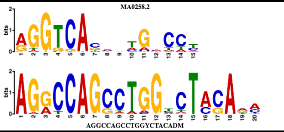 |
| | Pparg | 4.67E-04 | 6.69E-01 | 4.46E-01 | 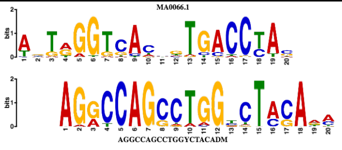 |
| | Sp2 | 2.02E-06 | 2.90E-03 | 1.91E-03 | 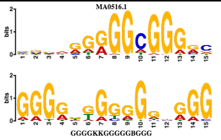 |
| | Rreb1 | 3.65E-06 | 5.23E-03 | 2.48E-03 | 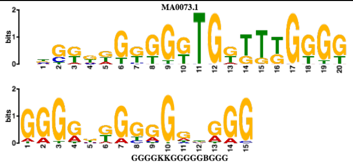 |
| | Egr1 | 1.28E-05 | 1.83E-02 | 4.04E-03 | 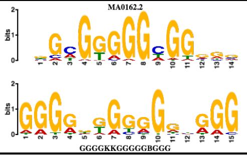 |
| | E2f3 | 2.17E-05 | 3.11E-02 | 5.60E-03 | 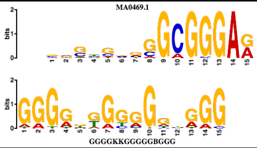 |
| | Sp4 | 4.48E-05 | 6.42E-02 | 1.06E-02 | 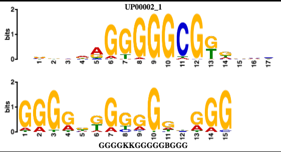 |
| | Egr2 | 6.18E-05 | 8.87E-02 | 1.35E-02 | 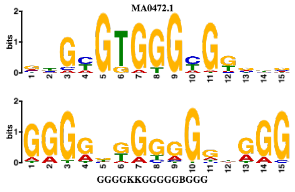 |
| | Sp3 | 1.15E-04 | 1.64E-01 | 2.33E-02 | 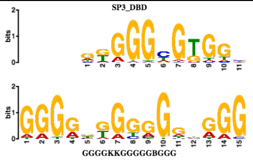 |
| | Sp1 | 1.49E-04 | 2.13E-01 | 2.82E-02 | 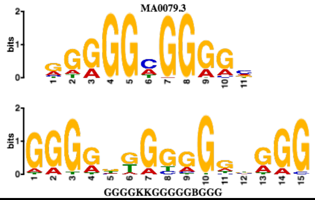 |

| | | | | | |
|---|---|---|---|---|---|
| | Smad3 | 3.01E-04 | 4.32E-01 | 4.98E-02 | 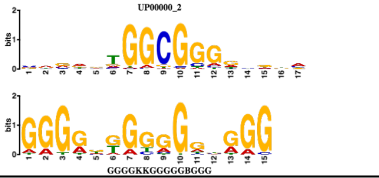  |
| | Klf5 | 7.01E-04 | 1.01E-01 | 4.88E-02 | 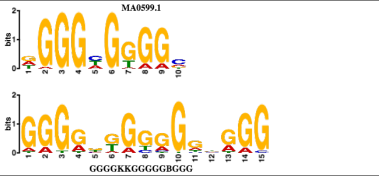  |
| | Stat1 | 1.73E-05 | 2.52E-01 | 1.20E-02 | 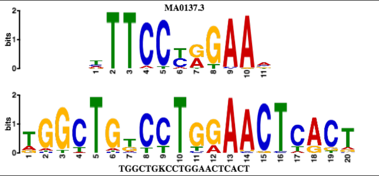  |
| | Rreb1 | 2.66E-08 | 3.81E-05 | 7.57E-05 | 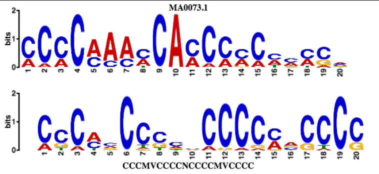  |
| | Zfp281 | 6.49E-08 | 9.30E-05 | 9.24E-05 | 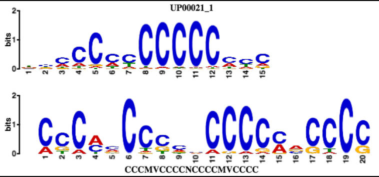  |
| | Zfp740 | 2.59E-07 | 3.71E-04 | 2.46E-04 | 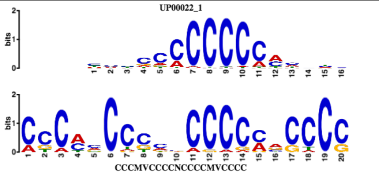  |
| | Nr5a2 | 2.60E-06 | 3.75E-03 | 7.49E-03 | 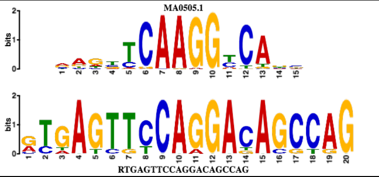  |
| | Bcl6 | 6.34E-05 | 9.09E-03 | 4.06E-02 | 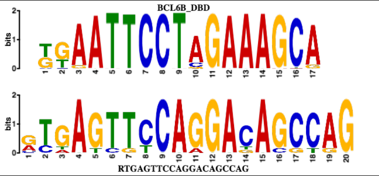  |
| | Hnf4a | 1.57E-04 | 2.25E-02 | 3.36E-02 | 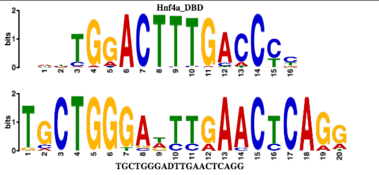  |
| | Ewsr1-Fli1 | 1.95E-06 | 2.79E-03 | 2.79E-03 | 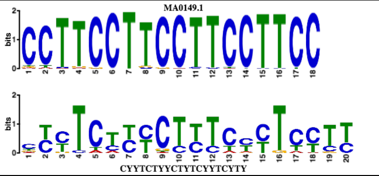  |
