## Supplementary Materials for "An unexpected contribution of lincRNA splicing to enhancer function"

**Supplementary Table ST4**

| | | | | |
|---|---|---|---|---|
| chr1 | 59362216 | 59362218 | ENSG00000232453.1 | rs3015311 |
| chr11 | 61514820 | 61514822 | ENSG00000124915.6 | rs4963390 |
| chr12 | 90676335 | 90676337 | ENSG00000258183.1 | rs2731262 |
| chr12 | 121830032 | 121830034 | ENSG00000258435.1 | rs7962143 |
| chr12 | 123199340 | 123199342 | ENSG00000256249.1 | rs140288839 |
| chr13 | 41040641 | 41040643 | ENSG00000215483.4 | rs17061303 |
| chr14 | 86537204 | 86537206 | ENSG00000258733.1 | rs113712266 |
| chr14 | 101324134 | 101324136 | ENSG00000214548.10 | rs116907618 |
| chr16 | 29224227 | 29224229 | ENSG00000260517.1 | rs74657328 |
| chr16 | 48778240 | 48778242 | ENSG00000260086.1 | rs78922375 |
| chr16 | 58467852 | 58467854 | ENSG00000260186.1 | rs4784938 |
| chr18 | 39792326 | 39792328 | ENSG00000267586.2 | rs8087600 |
| chr19 | 4470610 | 4470612 | ENSG00000267011.1 | rs10407114 |
| chr19 | 11755051 | 11755053 | ENSG00000197332.7 | rs113894173 |
| chr19 | 35897980 | 35897982 | ENSG00000205786.4 | rs35138704 |
| chr19 | 53703874 | 53703876 | ENSG00000269051.1 | rs17206931 |
| chr2 | 8718697 | 8718699 | ENSG00000236008.1 | rs10929538 |
| chr2 | 12698570 | 12698572 | XLOC_9410 | rs116218070 |
| chr2 | 12698570 | 12698572 | ENSG00000224184.1 | rs116218070 |
| chr2 | 59251968 | 59251970 | ENSG00000233723.3 | rs17049881 |
| chr2 | 70275820 | 70275822 | XLOC_10165 | rs72841112 |
| chr2 | 128146761 | 128146763 | ENSG00000236682.1 | rs2276683 |
| chr2 | 181968203 | 181968205 | ENSG00000238171.1 | rs75582374 |
| chr2 | 182187385 | 182187387 | ENSG00000234663.1 | rs13397327 |
| chr2 | 196397967 | 196397969 | ENSG00000223466.1 | rs16837006 |
| chr21 | 43444412 | 43444414 | ENSG00000237232.3 | rs220219 |
| chr22 | 36792160 | 36792162 | ENSG00000223695.1 | rs79674812 |
| chr3 | 98061273 | 98061275 | ENSG00000251088.1 | rs76255466 |
| chr4 | 119517064 | 119517066 | ENSG00000260404.2 | rs4001379 |
| chr4 | 185303530 | 185303532 | XLOC_13468 | rs1105754 |
| chr5 | 44808879 | 44808881 | XLOC_13971 | rs78665523 |
| chr6 | 2329566 | 2329568 | ENSG00000250903.4 | rs77773385 |
| chr6 | 2399097 | 2399099 | ENSG00000250903.4 | rs12210286 |
| chr7 | 35772457 | 35772459 | ENSG00000227544.4 | rs115661829 |
| chr7 | 135715781 | 135715783 | ENSG00000230649.2 | rs28625723 |
| chr8 | 2679913 | 2679915 | ENSG00000253853.1 | rs13265741 |
| chr8 | 130382550 | 130382552 | ENSG00000229140.4 | rs16904052 |
| chr9 | 35790150 | 35790152 | ENSG00000227388.2 | rs7866903 |

| SS_SNP_id | elincRNA_id | target_id |
|---|---|---|
| rs12210286 | ENSG00000250903.4 | ENSG00000112699.6 |
| rs7866903 | ENSG00000227388.2 | ENSG00000137078.4 |
| rs7866903 | ENSG00000227388.2 | ENSG00000137076.14 |
| rs7866903 | ENSG00000227388.2 | ENSG00000107185.8 |
| rs7866903 | ENSG00000227388.2 | ENSG00000159899.10 |
| rs7866903 | ENSG00000227388.2 | ENSG00000137133.6 |
| rs7866903 | ENSG00000227388.2 | ENSG00000070610.10 |
| rs7866903 | ENSG00000227388.2 | ENSG00000137098.9 |
| rs7866903 | ENSG00000227388.2 | ENSG00000107140.11 |
| rs7866903 | ENSG00000227388.2 | ENSG00000137103.12 |
| rs7866903 | ENSG00000227388.2 | ENSG00000107175.6 |
| rs35138704 | ENSG00000205786.4 | ENSG00000126262.4 |
| rs35138704 | ENSG00000205786.4 | ENSG00000105698.11 |
| rs35138704 | ENSG00000205786.4 | ENSG00000161249.16 |
| rs220219 | ENSG00000237232.3 | ENSG00000157617.12 |
| rs220219 | ENSG00000237232.3 | ENSG00000173276.9 |
| rs220219 | ENSG00000237232.3 | ENSG00000141956.9 |
