## Supplementary Materials for "An unexpected contribution of lincRNA splicing to enhancer function"

**Supplementary Table ST5**

| Cell Type | Protein of interest | Experiment type |
|---|---|---|
| mESC | Total cell | CAGE |
| NPC | Total cell | CAGE |
| Neuron | Total cell | CAGE |
| mESC | Total cell | HiC |
| NPC | Total cell | HiC |
| Neuron | Total cell | HiC |
| mESC | Total cell | RNA-seq |
| hESC | Total cell | RNA-seq |
| mESC | RNAPII | GRO-seq |
| mESC | RNAPII | ChIP-seq |
| mESC | RNAPII-S5P | ChIP-seq |
| mESC | RNAPII-S2P | ChIP-seq |
| mESC | H3K4me1 | ChIP-seq |
| mESC | H3K27ac | ChIP-seq |
| mESC | DHSI | DNase-seq |
| mESC | Cbp | ChIP-seq |
| mESC | p300 | ChIP-seq |
| mESC | Smad3 | ChIP-seq |
| mESC | Klf5 | ChIP-seq |
| mESC | Ctcf | ChIP-seq |
| mESC | Smc1 | ChIP-seq |
| mESC | Smc3 | ChIP-seq |
| mESC | Nipbl | ChIP-seq |
| mESC | Med1 | ChIP-seq |
| mESC | Med12 | ChIP-seq |
| Population LCL | Total cell | Genotyping |
| Population LCL | Total cell | RNA-seq |
| LCL (GM12878) | Total cell | HiC |
| LCL (GM12878) | Total cell | CAGE |
| LCL (GM12878) | Total cell | RNA-seq |

| GEO accession | Reference | |
|---|---|---|
| GSE59027 | Fraser et al 2016 | |
| GSE59027 | Fraser et al 2016 | |
| GSE59027 | Fraser et al 2016 | |
| GSE59027 | Fraser et al 2016 | |
| GSE59027 | Fraser et al 2016 | |
| GSE59027 | Fraser et al 2016 | |
| GSE58757 | Tan et al 2015 | |
| ENCODE | Encode Project Consortium 2012 | |
| GSE27037 | Min et al 2011 | |
| ENCODE | Encode Project Consortium 2012 | |
| GSE34520 | Brookes et al 2012 | |
| GSE34520 | Brookes et al 2012 | |
| ENCODE | Encode Project Consortium 2012 | |
| ENCODE | Encode Project Consortium 2012 | |
| ENCODE | Encode Project Consortium 2012 | |
| GSE51522 | Hnisz et al 2013 | |
| ENCODE | Chen et al 2008 | ? |
| GSE11431 | Chen et al 2008 | |
| GSE49848 | Aksoy et al 2014 | |
| ENCODE | Encode Project Consortium 2012 | |
| GSE22557 | Kagey et al 2010 | |
| GSE22557 | Kagey et al 2010 | |
| GSE22557 | Kagey et al 2010 | |
| GSE22557 | Kagey et al 2010 | |
| GSE22557 | Kagey et al 2010 | |
| 1000 Genomes | 1000 Genomes Project Consortium 2012 | |
| Geuvadis | Lappalainen et al 2013 | |
| GSE63525 | Rao et al 2014 | |
| Fantom | Hon et al 2016 | |
| Fantom | Hon et al 2016 | |
