## Supplementary Materials for "An unexpected contribution of lincRNA splicing to enhancer function"

#### SUPPLEMENTARY FIGURE LEGENDS

**Figure S1. elincRNA splicing is conserved and efficient.** (A) Distribution of the GC-content of exons and introns of single- and multi-exonic elincRNAs (red), other expressed lincRNAs (blue), protein-coding genes (green) and their respective flanking regions (grey). Distribution of the max entropy scores of (B) 5' and (C) 3' splice sites (SS) for multi-exonic elincRNAs (red), other expressed lincRNAs (blue), and protein-coding genes (green). (D) Distribution of the average synthesis and degradation rates for elincRNAs (red), other expressed lincRNAs (blue) and protein-coding genes (green). (E) Mouse-human pairwise substitution rate for exons of multi-exonic (red) and single-exonic (grey) elincRNAs, other expressed lincRNAs (blue), protein-coding genes (green), and their nearby ancestral repeats (ARs, black). Substitution rates relative to proximal ARs are shown in figure insets. (F) Distribution of the RNA synthesis rates of multi-exonic elincRNAs (red), other expressed lincRNAs (blue) and protein-coding genes (green), as well as their single-exonic counterparts (grey). (G) Metagene plot of mESCs GRO-seq reads centered at transcription initiation regions (TIRs) of multi-exonic (red) and single-exonic (grey) elincRNAs and eRNAs (yellow). (H) Distribution of RNAPII travelling ratio (TR) for multi-exonic (red) and single-exonic (grey) elincRNAs, eRNAs (yellow), other expressed lincRNAs (blue) and protein-coding genes (green). Metagene plots and distribution (figure insets) of ChIP-seq reads for RNAPII with (I) phosphorylated-Serine 5 (S5P) and (J) phosphorylated Serine 2 (S2P) at their C-terminal domain centered at TIRs of multi-exonic (red) and single-exonic (grey) elincRNAs and eRNAs (yellow). Differences between groups were tested using a two-tailed Mann-Whitney *U* test.

**Figure S2. Multi-exonic elincRNAs are associated with higher enhancer activity.** Distribution of the fold difference (FD) in transcription (measured as CAGE TPM) of the closest gene that is expressed in the same embryonic neurogenesis stage as (A) elincRNAs (red), other mESC-expressed lincRNAs (oth-lincRNAs, blue) and protein-coding genes (PCGs, green) and (B) multi-exonic elincRNAs (red), other expressed lincRNAs (blue) and protein-coding genes (green) compared to their single-exonic counterparts (grey). Fold difference of neighboring gene transcription is calculated between the two cellular stages across neuronal differentiation, where the expression level of the reference locus (elincRNA, oth-lincRNA, or PCG) is maximal and minimal. Metagene plots and distribution (figure insets) of (C) p300, (D) Smad3, and (E) Klf5 ChIP-seq reads in mESCs at the transcription initiation regions of multi-exonic (red) and single-exonic (grey) elincRNAs, as well as eRNAs (yellow). Differences between groups were tested using a two-tailed Mann-Whitney *U* test. * $p < 0.05$; ** $p < 0.01$.

**Figure S3. Multi-exonic elincRNAs are associated with modulation of local chromosomal architecture.** Metagene plots and distribution (figure insets) of (A) Ctcf, (B) Smc1, (C) Smc3, (D) Nipbl, (E) Med1 and (E) Med12 ChIP-seq reads in mESCs at transcription initiation regions of multi-exonic (red) and single-exonic (grey) elincRNAs, as well as eRNAs (yellow). Differences between groups were tested using a two-tailed Mann-Whitney *U* test. * $p < 0.05$; ** $p < 0.01$, *** $p < 0.001$.

**Figure S4. Multi-exonic elincRNAs are associated with cell-type specific TAD boundaries.** (A) Distribution of the distance between single-exonic elincRNA (grey) and eRNA (yellow) transcription initiation site to their nearest TAD border in mESCs, neuronal precursor cells (NPCs) and neurons. Metagene plots of CAGE reads centered at enhancers that transcribe (B) multi-exonic elincRNAs and (C) eRNAs and located at TAD boundaries that are either cell-stage invariant (conserved) or specific (non-conserved) across embryonic neurogenesis (mESC to NPC to Neuron). Sense (red) and antisense (blue) reads denote those that map to the same or opposite strand, respectively, as the direction of their cognate TIRs.

**Figure S5. Impact of elincRNA splicing on *cis*-gene regulation in the human population.** (A) Joint QTL variants (joint seQTLs/esQTLs, yellow box), which are SNPs that are associated with both elincRNA (red boxes) splicing (sQTLs) and the expression levels (eQTLs) of their putative *cis*-target genes (green boxes), were used to test the likelihood that target-eQTL associations are indirect effect (dotted arrow) of elincRNA-sQTL associations (solid arrow)

(Model 1, forward model). Conversely, the reverse model (Model 2) tests the probability that target-eQTLs (solid arrow) mediates elincRNA-sQTL associations (dotted arrow). (B) The proportion of all elincRNA-sQTLs that are jointly associated with the expression levels (eQTLs) of their putative target protein-coding genes (joint seQTLs) (red) out of all elincRNA-sQTLs (grey) [Forward model] compared to the proportion of all target-eQTLs that are jointly associated with elincRNA-sQTLs (joint esQTLs, green) out of all target-eQTLs (grey) [Reverse model]. Differences between groups were tested using a two-tailed Fisher's exact test. (C) Distribution of causal mediation testing (Sobel's test) adjusted p-values (Benjamini-Hochberg multiple testing correction, qval) associated with the forward and reverse models (as illustrated in A) for elincRNAs. Dotted red lines denote significance threshold at qval < 0.05.

#### SUPPLEMENTARY TABLE LEGENDS

**Table ST1.** Loci of single- and multi-exonic mESC elincRNAs (mm9) and their respective number of exons.

**Table ST2.** Metabolic rates of elincRNAs, including their rate of synthesis, processing and degradation.

**Table ST3.** Motifs of transcription factor binding sites enriched at transcription initiation regions of multi-exonic elincRNAs.

**Table ST4.** Single nucleotide polymorphisms (SNPs) at multi-exonic elincRNA splice sites.

**Table ST5.** Publically available datasets used in the analysis.

**Supp Figure S1.**

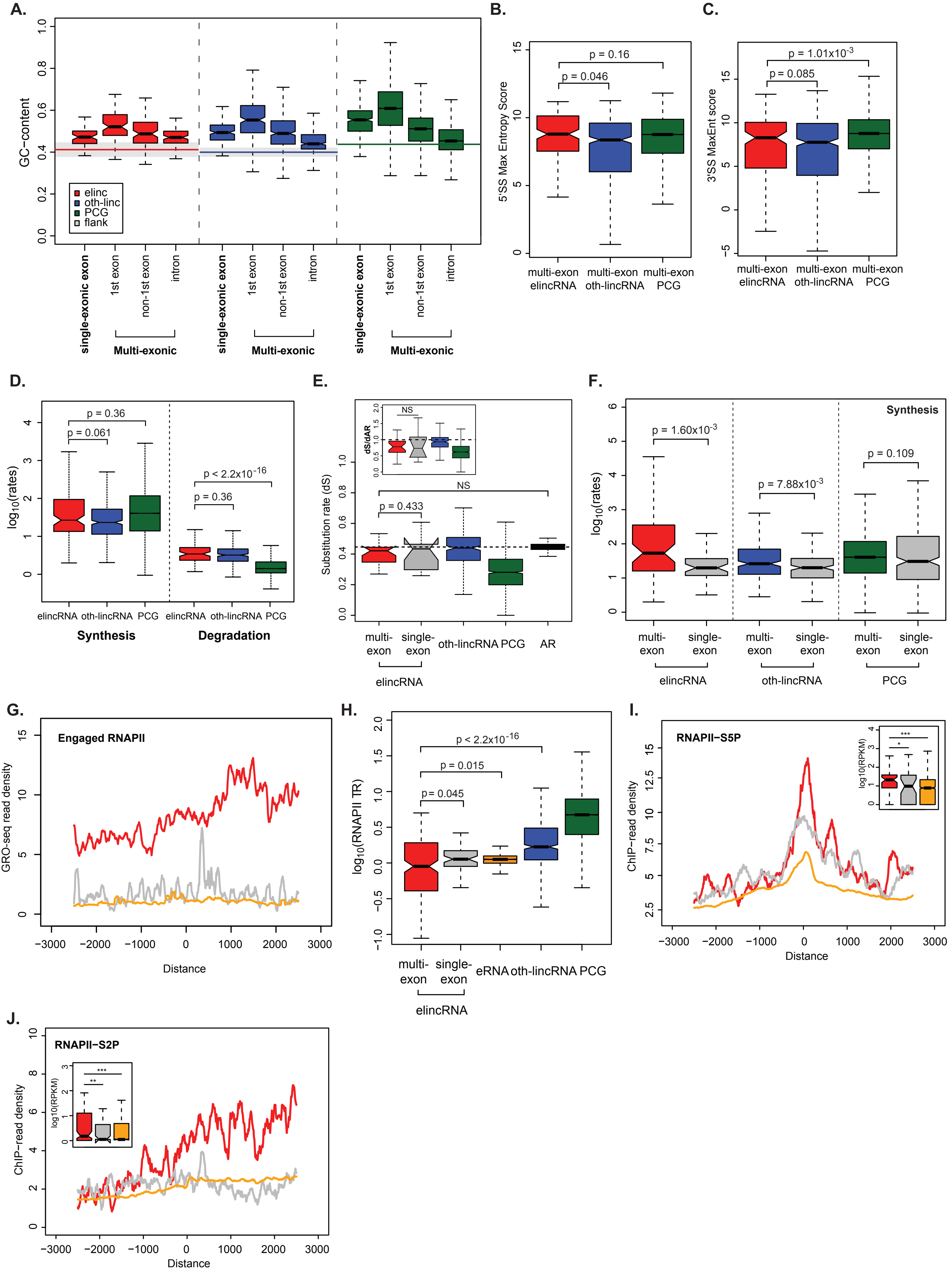

### Supp Figure S2.

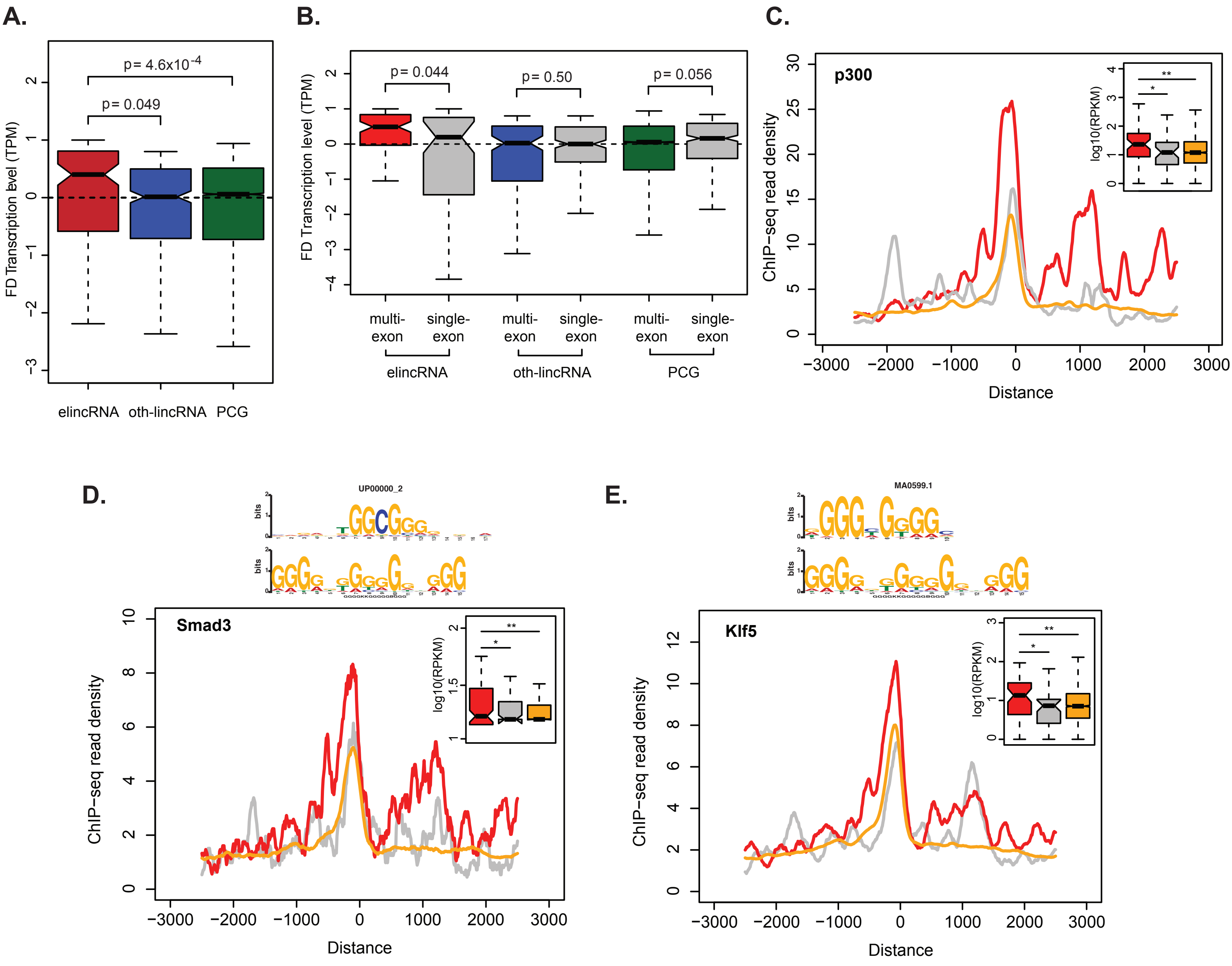

### Supp Figure S3.

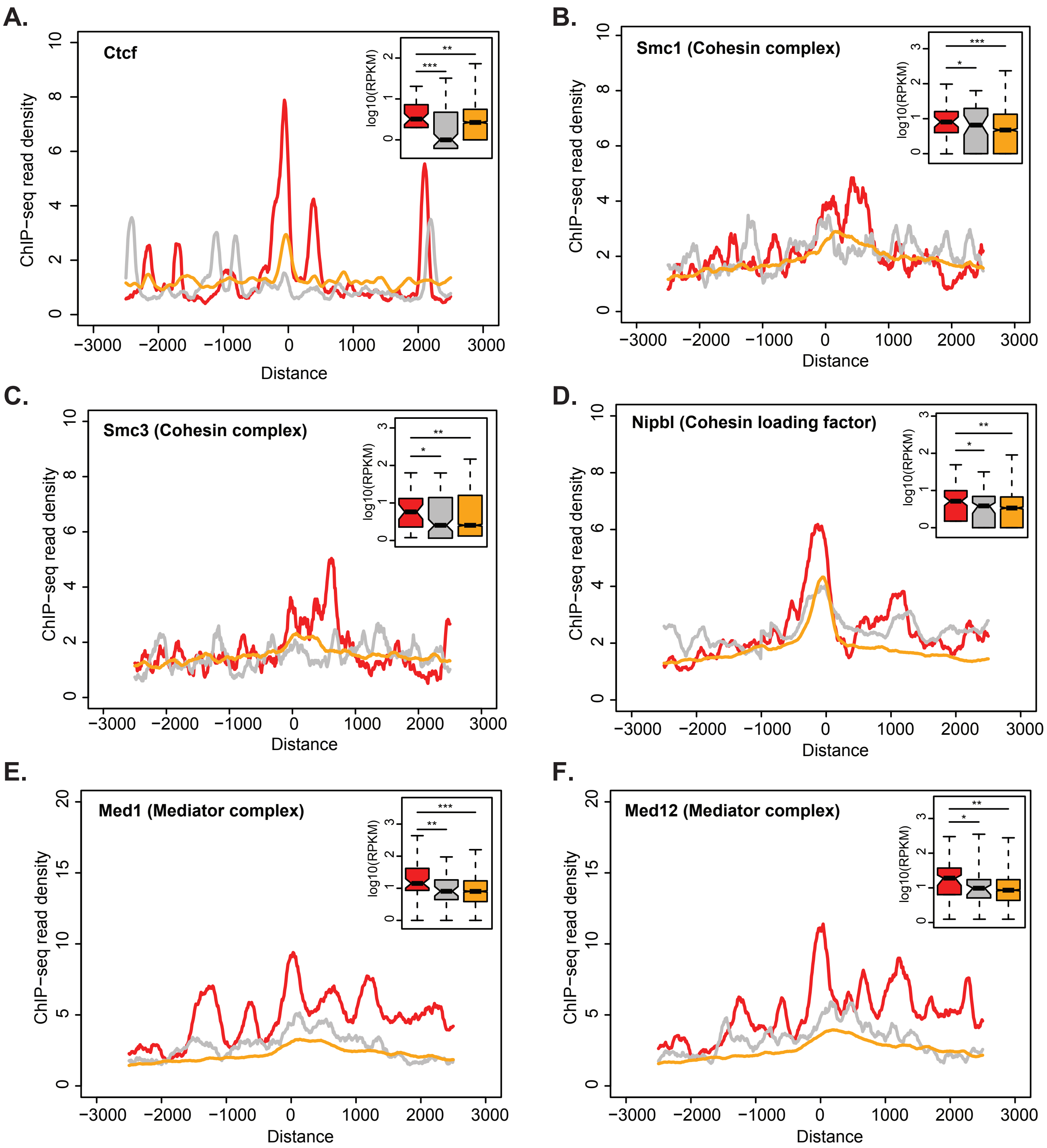

#### Supp Figure S4.

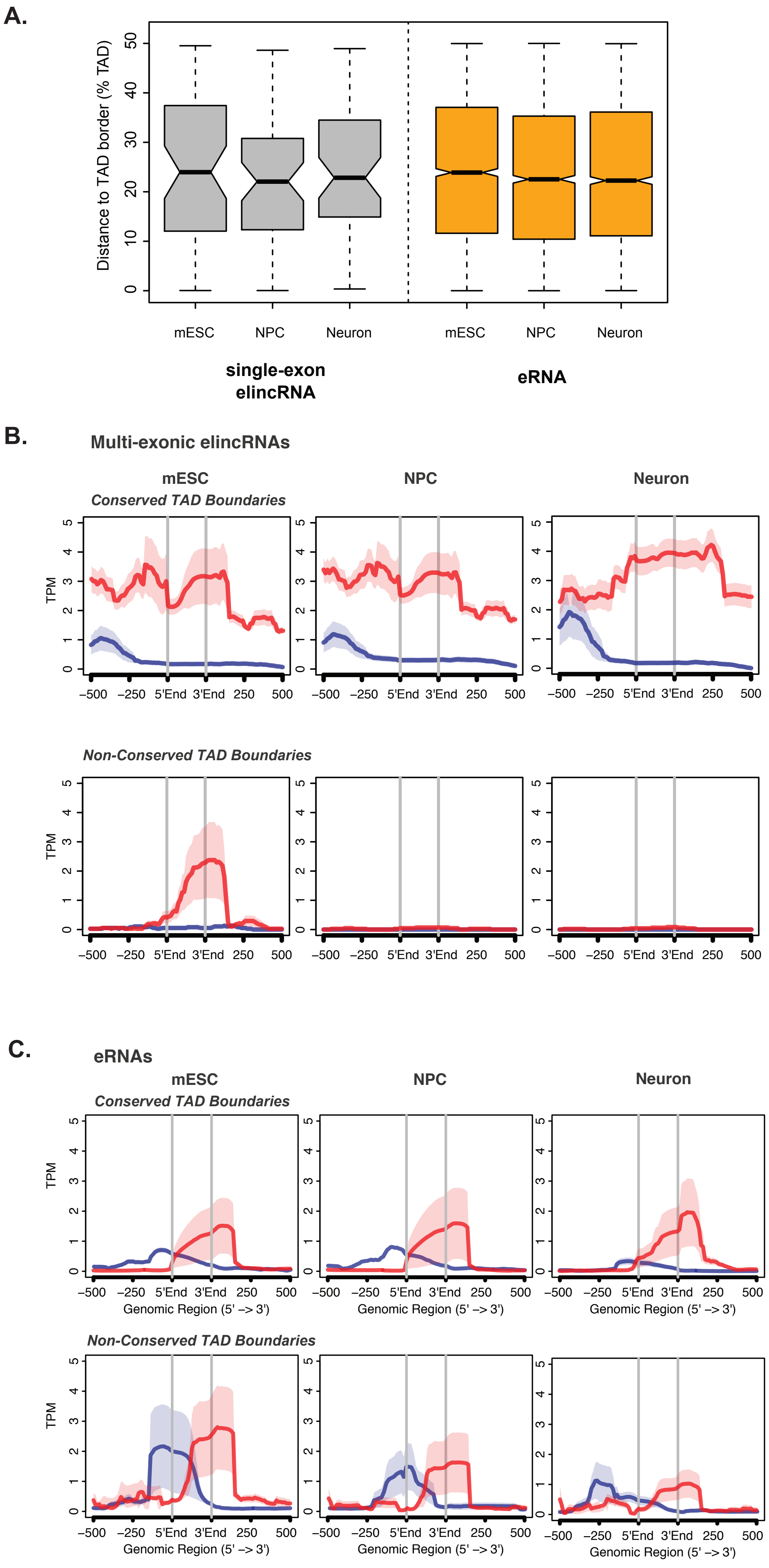

#### Figure S5.

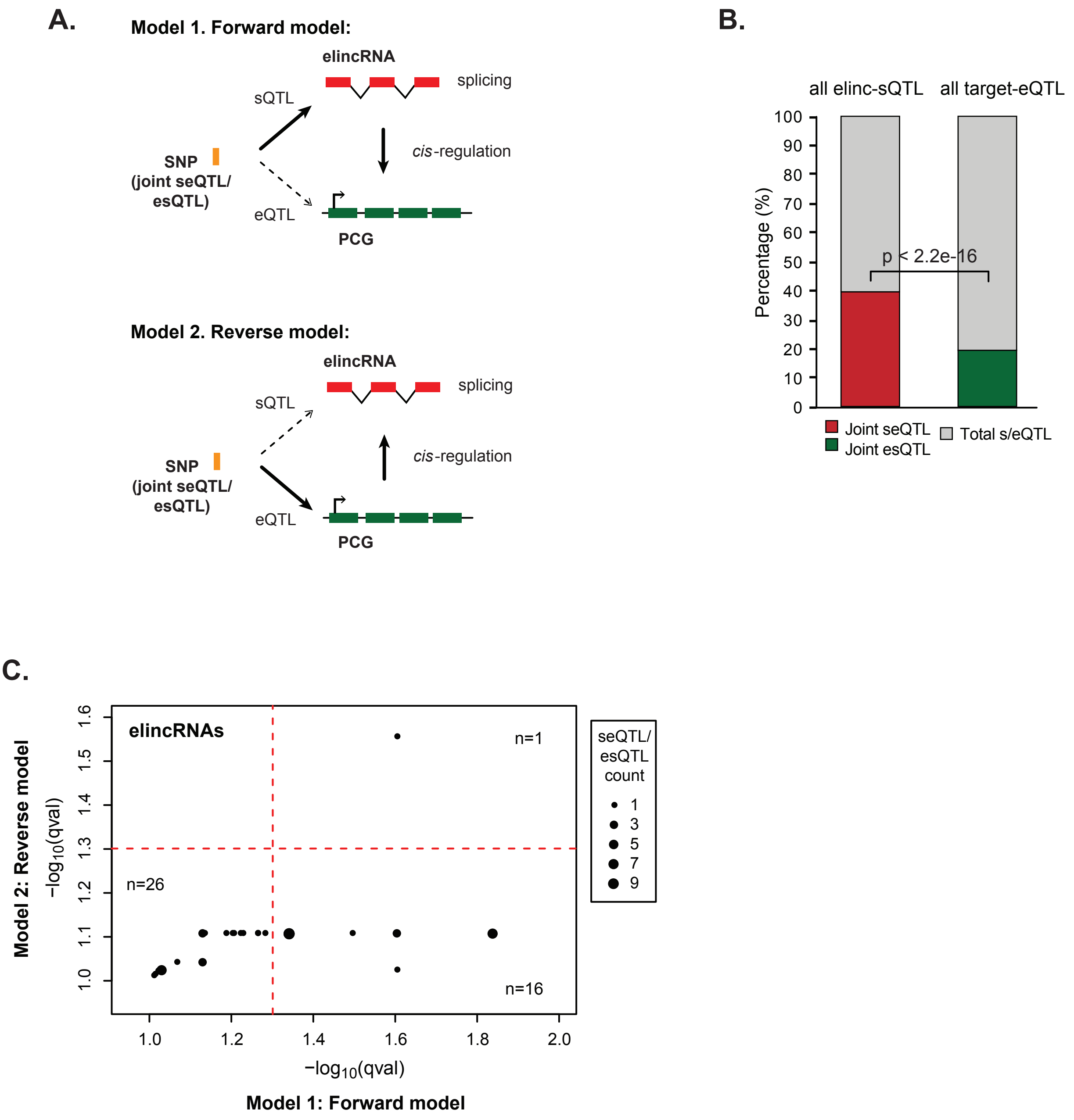
A.
Model 1. Forward model:
elincRNA
splicing
sQTL
cis-regulation
SNP
(joint seQTL/
esQTL)
eQTL
PCG
Model 2. Reverse model:
elincRNA
splicing
sQTL
cis-regulation
SNP
(joint seQTL/
esQTL)
eQTL
PCG
B.
all elinc-sQTL
all target-eQTL
Percentage (%)
p < 2.2e-16
Joint seQTL
Joint esQTL
Total s/eQTL
C.
elincRNAs
n=1
n=26
n=16
Model 2: Reverse model
−log10(qval)
−log10(qval)
Model 1: Forward model
seQTL/
esQTL
count
1
3
5
7
9
